# Multivariate pattern similarity of hemodynamic brain responses to spoken and written words increases during primary school

**DOI:** 10.64898/2026.05.08.723790

**Authors:** Kazuki Maruo, Roman Kessler, Falk Huettig, Michael A. Skeide

**Affiliations:** Institute of Child and Adolescent Psychiatry, Kiel University and University Medical Center Schleswig–Holstein, Niemannsweg 147, 24105 Kiel, Germany; The Cultural Brain Research Group, Max Planck Institute for Psycholinguistics, Wundtlaan 1, Nijmegen 6525, Netherlands

## Abstract

Learning to read requires binding auditory and visual information together, yet how the developing brain maps information across sensory modalities at different levels of reading experience remains poorly understood. To shed light on this topic we employed functional magnetic resonance imaging to investigate hemodynamic brain responses during spoken and written word recognition in primary school children learning to read the alpha-syllabic Hindi Devanagari script in the first, second, or fourth grade of primary school. Multivariate audiovisual similarity of activation patterns in the inferior frontal gyrus, inferior parietal lobule, posterior superior and middle temporal gyrus, and temporo-occipital cortex, increased with school grade. This effect was independent from the behavioral performance in a semantic judgement task performed in the scanner. Our results suggest that, as children learn to read during primary school, their brain responses to written language become more similar to their brain responses to spoken language.

## Introduction

Learning to read is a fundamental enculturation process in formal education, creating a versatile cognitive tool for lifelong learning. During reading instruction, children learn to map written letters or characters onto spoken sounds to form links between written word forms, their corresponding spoken word forms, and word meanings, a process known as orthographic mapping (Ehri, 2014; Miles & Ehri, 2019; Schmalz et al., 2020; Share, 1995). Difficulties in linking visual and auditory information have been consistently identified in children diagnosed with developmental dyslexia (Blau et al., 2010; Blomert, 2011; Blomert & Willems, 2010; Castles et al., 2018; Fraga González et al., 2017; Golden & Zenhausern, 1983).

Converging evidence from functional magnetic resonance imaging (fMRI) experiments on auditory language comprehension suggests that left inferior frontal, inferior parietal, superior temporal, and middle temporal cortices are involved in phonological and semantic information processing from early childhood onwards (Enge et al., 2020; Friederici, 2011; Malik-Moraleda et al., 2022; Skeide & Friederici, 2016; Turker et al., 2023; Vigneau et al., 2006). Similarly, a recent meta-analysis of written language comprehension in adults also identified large left-hemispheric activation clusters in the inferior frontal, inferior parietal, and posterior superior and middle temporal regions (Turker et al., 2025). Furthermore, there is evidence from adults across four different languages that hemodynamic responses to spoken and written words overlap in these regions (Rueckl et al., 2015). These findings indicate that left frontal, parietal and temporal regions are capable of processing language independently of the sensory input modality (Fedorenko et al., 2024).

Developmental fMRI work indicates that overlapping hemodynamic responses to spoken and written words can already be detected in the inferior frontal, inferior parietal, and superior temporal cortices of beginning readers (Chyl et al., 2018; Dębska et al., 2021; Dębska et al., 2026; McNorgan et al., 2014; Preston et al., 2016). Moreover, a multivariate analysis targeting distributed patterns of hemodynamic activity demonstrated that the representational similarity of responses to spoken and logographic written Chinese words is larger in adult college students compared to fifth graders within left frontal and temporal perisylvian regions (Yan et al., 2024). It remains unknown, however, whether such differences in audiovisual similarity already emerge during earlier stages of formal reading instruction and whether they generalize to other writing systems.

To close these current research gaps, we presented words and pseudowords either in the auditory or visual sensory modality to first, second, and fourth graders while collecting fMRI data. In the scanner, children performed a semantic judgement task in which they had to answer whether they had seen or heard a word corresponding to an animal or not. The sole purpose of this task was to maintain attention and motivation during scanning. Pseudowords were contrasted against words since they share comparable orthographic properties while lacking meaningful lexical representations. Going beyond alphabetic scripts like English and logographic scripts like Chinese, the focus of our study was on children learning the alpha-syllabic Devanagari script, one of the world’s most widely used but so far understudied writing systems. To quantify the similarity of hemodynamic responses between spoken and written words, we performed a region-of-interest based correlational analysis in twelve well-established language regions in the left hemisphere (Fig. 1), including inferior frontal, inferior parietal, superior temporal, middle temporal, and fusiform cortices, (Allison et al., 1994; Cohen et al., 2002; Dehaene et al., 2010; Enge et al., 2020; Friederici, 2011; Malik-Moraleda et al., 2022; Skeide & Friederici, 2016; Turker et al., 2023; Turker et al., 2025). Specifically, we tested whether audiovisual similarity differed across school grades and whether cross-modal differences were dissociable from unimodal differences of spoken or written word recognition.

**Figure 1.**
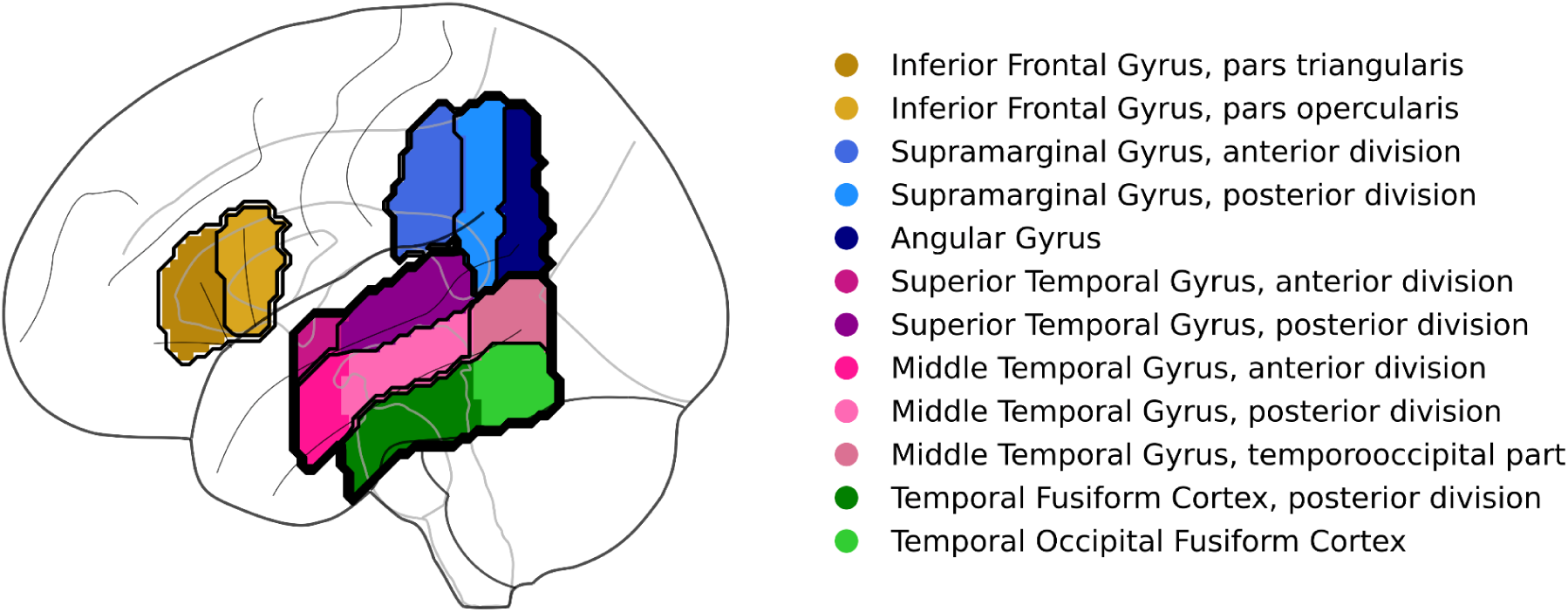
Regions of interest. The brain map displays twelve regions of interest in the left hemisphere selected from the Harvard–Oxford Cortical Atlas that have been associated with auditory and visual language comprehension in the literature.

We formulated the hypothesis that the left posterior superior temporal gyrus would exhibit an increase in audiovisual similarity across school grades. This hypothesis was based on converging findings reported in a recent meta-analysis relating this region to the integration of auditory and visual language information in adults (Gao et al., 2024). In addition, this hypothesis was grounded on developmental evidence for an increasing involvement of this region in audiovisual integration from preschool to school age (Karipidis et al., 2021). Finally, this hypothesis was motivated by a recent study in which this region revealed higher audiovisual similarity during word recognition in adult college students compared to children in fifth grade (Yan et al., 2024).

## Results

Demographic details, fMRI data quality indices and behavioral performance metrics are reported in Table 1.

**Table 1.** Sample characteristics.

| Variable | 1st grade (n = 18) | 2nd grade (n = 19) | 4th grade (n = 24) | Total (n = 61) |
| --- | --- | --- | --- | --- |
| Age, years | 6.3 ± 0.6 | 7.0 ± 0.7 | 9.1 ± 0.7 | 7.6 ± 1.4 |
| Sex, n (%) |  |  |  |  |
| Female | 9 (50.0%) | 9 (47.4%) | 12 (50.0%) | 30 (49.2%) |
| Male | 9 (50.0%) | 10 (52.6%) | 12 (50.0%) | 31 (50.8%) |
| Usable runs per subject, n | 5.2 ± 1.2 | 4.8 ± 1.2 | 5.7 ± 1.5 | 5.2 ± 1.3 |
| Task response rate per run, % | 90.7 ± 10.9 | 87.3 ± 13.7 | 92.2 ± 12.6 | 90.4 ± 12.6 |
| Task accuracy per run, % | 60.9 ± 16.8 | 59.6 ± 15.5 | 63.3 ± 18.8 | 61.5 ± 17.4 |
| Task reaction time per run, seconds | 2.2 ± 0.5 | 2.4 ± 0.3 | 2.0 ± 0.5 | 2.2 ± 0.4 |
| Average FD per run, mm<sup>a</sup> | 0.40 ± 0.26 | 0.39 ± 0.24 | 0.30 ± 0.24 | 0.35 ± 0.25 |

<sup>a</sup>FD, framewise displacement. mm, millimeters.

### Cross-modal audiovisual similarity and unimodal visual similarity differs across school grades

We first examined whether cross-modal audiovisual similarity within each region of interest (ROI) differed across school grades. To this end, similarity was computed as the spatial correlation between effect maps of hemodynamic responses to spoken and written words. We conducted a two-way mixed analysis of variance (ANOVA) with school grade as a between-subjects factor and ROI as a within-subjects factor. This test revealed a significant main effect of school grade (*F*(2, 58) = 7.11, *p* < .01, η_p_² = .19), a significant main effect of ROI (*F*(11, 638) = 17.21, *p* < .001, η_p_² = .23), and a significant interaction effect (*F*(22, 638) = 2.25, *p* < .01, η_p_² = .07). Regarding the main effect of ROI, post-hoc pairwise *t*-tests indicated that the posterior superior temporal gyrus (pSTG) exhibited significantly higher similarity than 10 of the 11 other ROIs (2.97 < *t*(60) < 8.84, *p_FDR_* < .01, 0.26 < Hedges’ g < 1.35), with the exception of the posterior supramarginal gyrus (pSMG) (*t*(60) = 1.33, *p_FDR_* = .24, Hedges’ g = 0.11).

Figure 2 illustrates post-hoc pairwise comparisons within each ROI, indicating significant differences between fourth graders and both first and second graders in the pars opercularis of the inferior frontal gyrus, the anterior and posterior division of the supramarginal gyrus, the temporo-occipital part of the middle temporal gyrus, and the temporo-occipital fusiform cortex (TOFC) (2.48 < *t* < 3.76, *p_FDR_* < .05, 0.74 < Hedges’ g < 1.10). Additionally, significant differences were identified between fourth graders and first graders in the pars triangularis of the inferior frontal gyrus (*t* = 2.60, *p_FDR_* < .05, Hedges’ g = 0.77), and between fourth graders and second graders in the aSTG and pSTG (2.62 < *t* < 2.63, *p_FDR_* < .05, 0.76 < Hedges’ g < 0.79). No statistically significant differences between first and second graders were observed in any of the ROIs.

**Figure 2.**
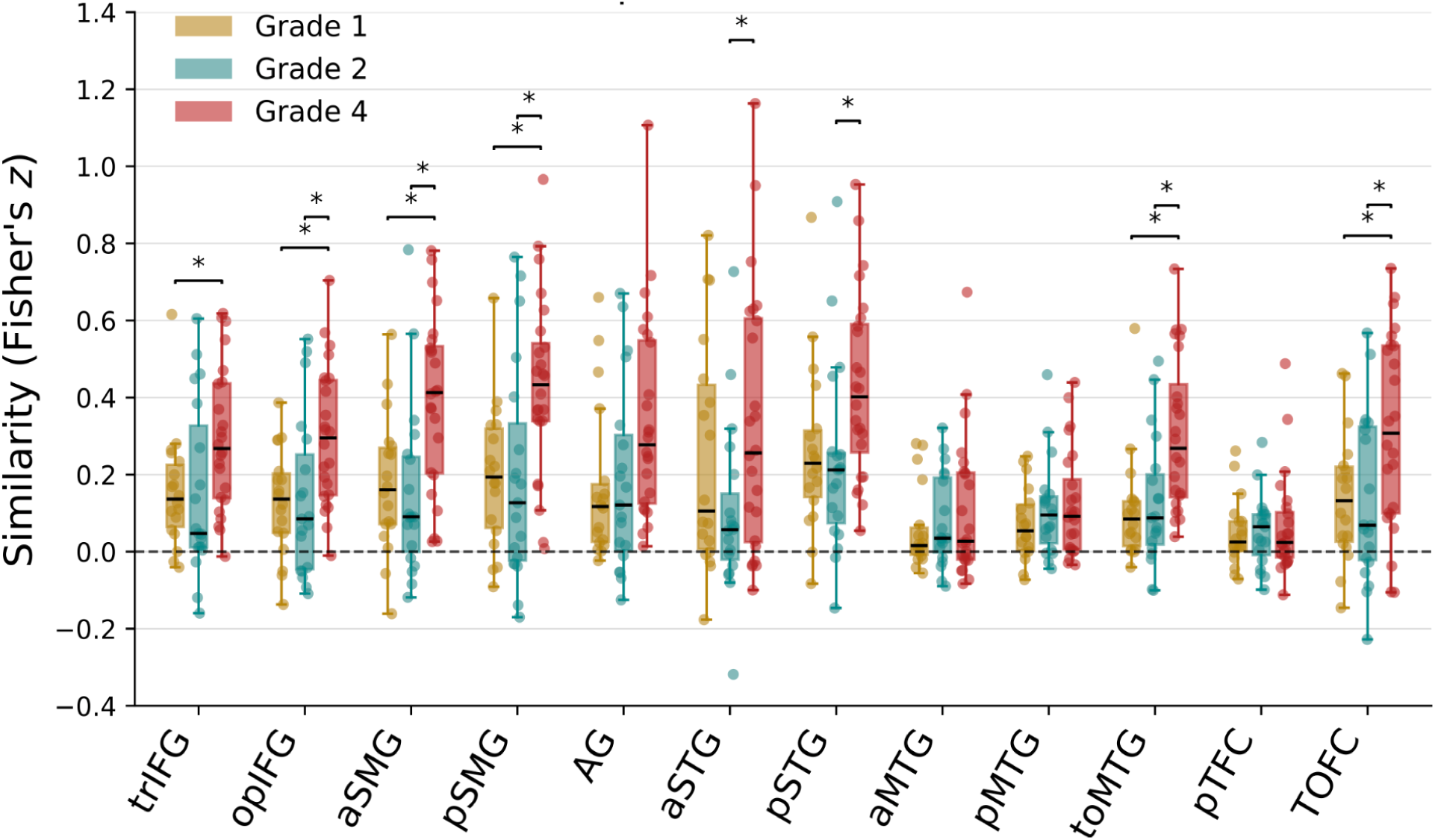
Cross-modal audiovisual similarity across regions of interest and school grades. Box plots illustrate audiovisual similarity (y-axes) across regions of interest (x-axes) and school grades. Each data point represents the estimated similarity value of each individual participant. Boxplots display the median and interquartile range (IQR), with whiskers extending to the most extreme values within 1.5 times the IQR from the first and third quartiles. Asterisks denote statistical significance (*p_FDR_* < .05) derived from post-hoc pairwise comparisons. trIFG, inferior frontal gyrus, pars triangularis; opIFG, inferior frontal gyrus, pars opercularis; aSMG, supramarginal gyrus, anterior division; pSMG, supramarginal gyrus, posterior division; AG, angular gyrus; aSTG, superior temporal gyrus, anterior division; pSTG, superior temporal gyrus, posterior division; aMTG, middle temporal gyrus, anterior division; pMTG, middle temporal gyrus, posterior division; toMTG, middle temporal gyrus, temporooccipital part; pTFC, temporal fusiform cortex, posterior division; TOFC, temporal occipital fusiform cortex.

A significant grade x ROI interaction effect was found for unimodal visual similarity (*F*(22, 638) = 1.99, *p* < .05, η_p_² = .064) and post-hoc comparisons were largely consistent with the cross-modal results, with the exception that no significant difference was found in the TOFC (Fig. 3). The interaction effect was also significant for unimodal auditory similarity (*F*(22, 638) = 1.64, *p* < .05, η_p_² = .054), but post-hoc tests revealed no significant pairwise differences (Fig. 3). Cross-modal audiovisual similarity of semantic information processing, estimated from the contrast between words and pseudowords, revealed no significant main effects of school grade (*F*(2, 58)=1.48, *p* = .237, η_p_² =.048) and ROI (*F*(11, 638) = 1.66, *p* = .10, η_p_² = .02), and no interaction effect (*F*(22, 638) = 0.74, *p* = .804, η_p_² = .025).

**Figure 3.**
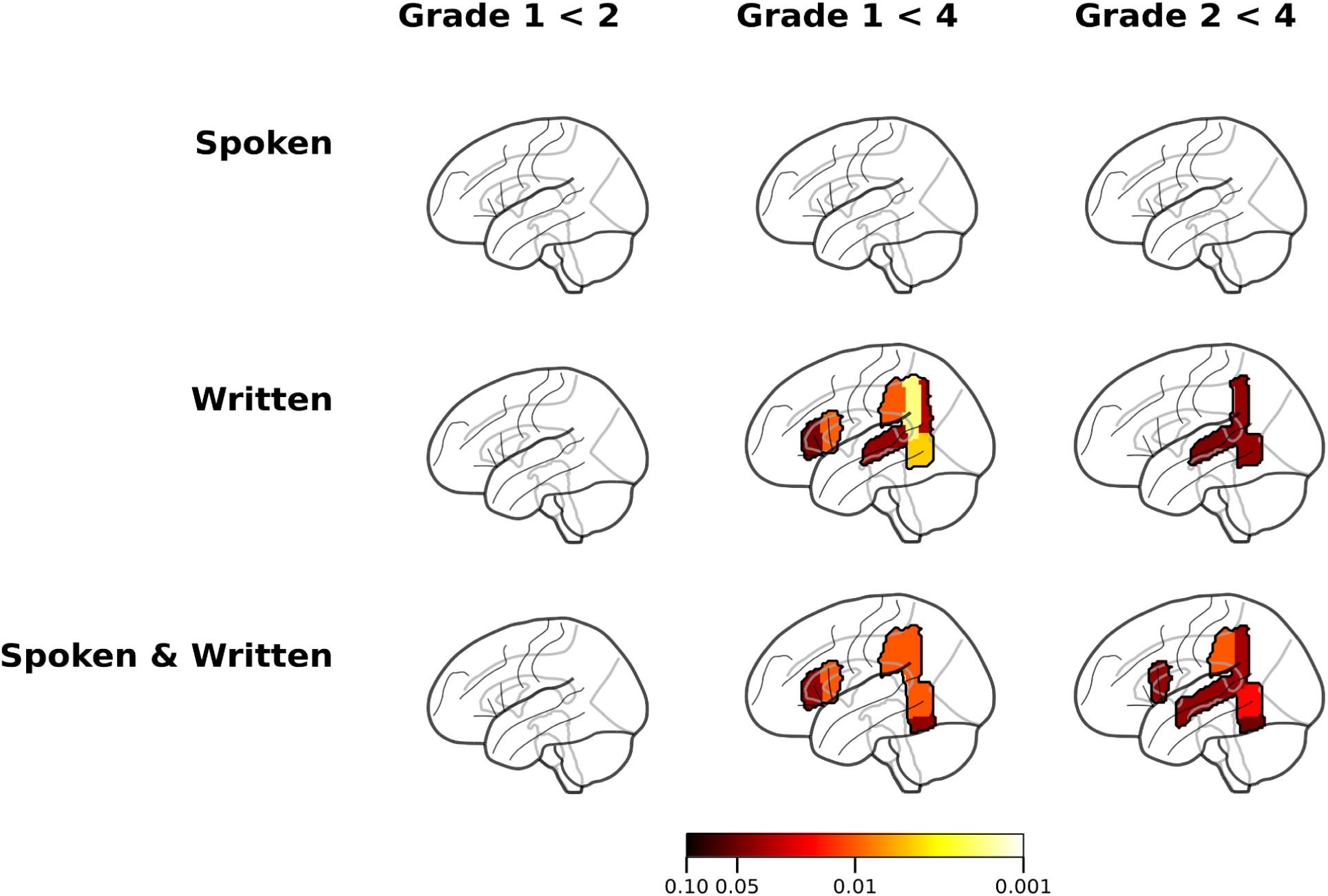
Regions of interest exhibiting significant differences across sensory modalities and school grades. The brain maps illustrate regions showing significant differences at a level of *p_FDR_* < .05 with respect to the similarity of hemodynamic responses to spoken words only (unimodal auditory similarity, top row), written words only (unimodal visual similarity, mid row), and both spoken and written words (cross-modal audiovisual similarity, bottom row). The color bar depicts false-discovery-rate-corrected *p*-values. *P*-values up to *p_FDR_* < .10 are also shown to report trends towards significance, but no such trends were observed.

### Cross-modal audiovisual similarity increases with school grade

To determine whether cross-modal and within-modal representational similarity increased in the course of primary school, we performed multiple linear regression analyses for each ROI. In these analyses, we treated school grade as the independent variable and included head motion (mean framewise displacement), sex, and behavioral performance in the semantic judgement task (mean reaction time) as covariates of no interest in the model. ROIs with significant positive regression coefficients for the effect of school grade are summarized in Table 1. Cross-modal audiovisual similarity increased with school grade in the left opIFG, aSMG, pSMG, pSTG, toMTG, and TOFC (0.04 < *β* < 0.07, 2.45 < *t* < 3.68, *pFDR* < .05). Except for the TOFC, a comparable developmental increase was observed for unimodal visual similarity (0.04 < *β* < 0.08, 2.47 < *t* < 4.24, *pFDR* < .05). Significant positive associations between school grade and unimodal auditory similarity were restricted to the pSMG and toMTG (0.07 < *β* < 0.08, 2.93 < *t* < 3.13, *pFDR* < .05). Following our hypothesis, Figure 4 illustrates cross-modal and unimodal similarity values, derived from the fitted regression models, in the pSTG. Audiovisual similarity also revealed a significant positive correlation with audiovisual coactivation indices derived from conventional voxel-level conjunction analysis (see Supplementary Information).

**Table 1.**
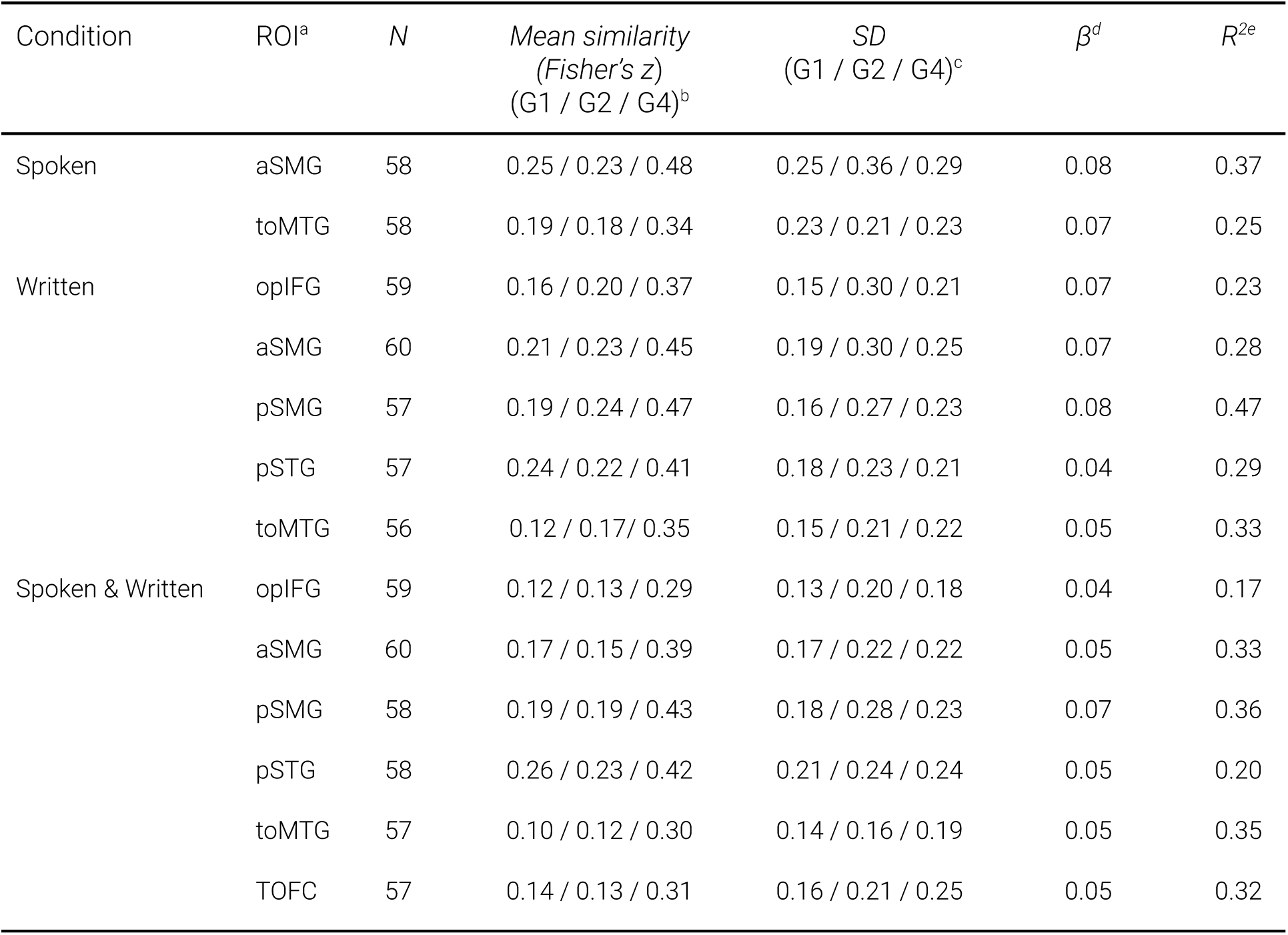
Significant associations between school grade and cross-modal or unimodal similarity.

**Figure 4.**
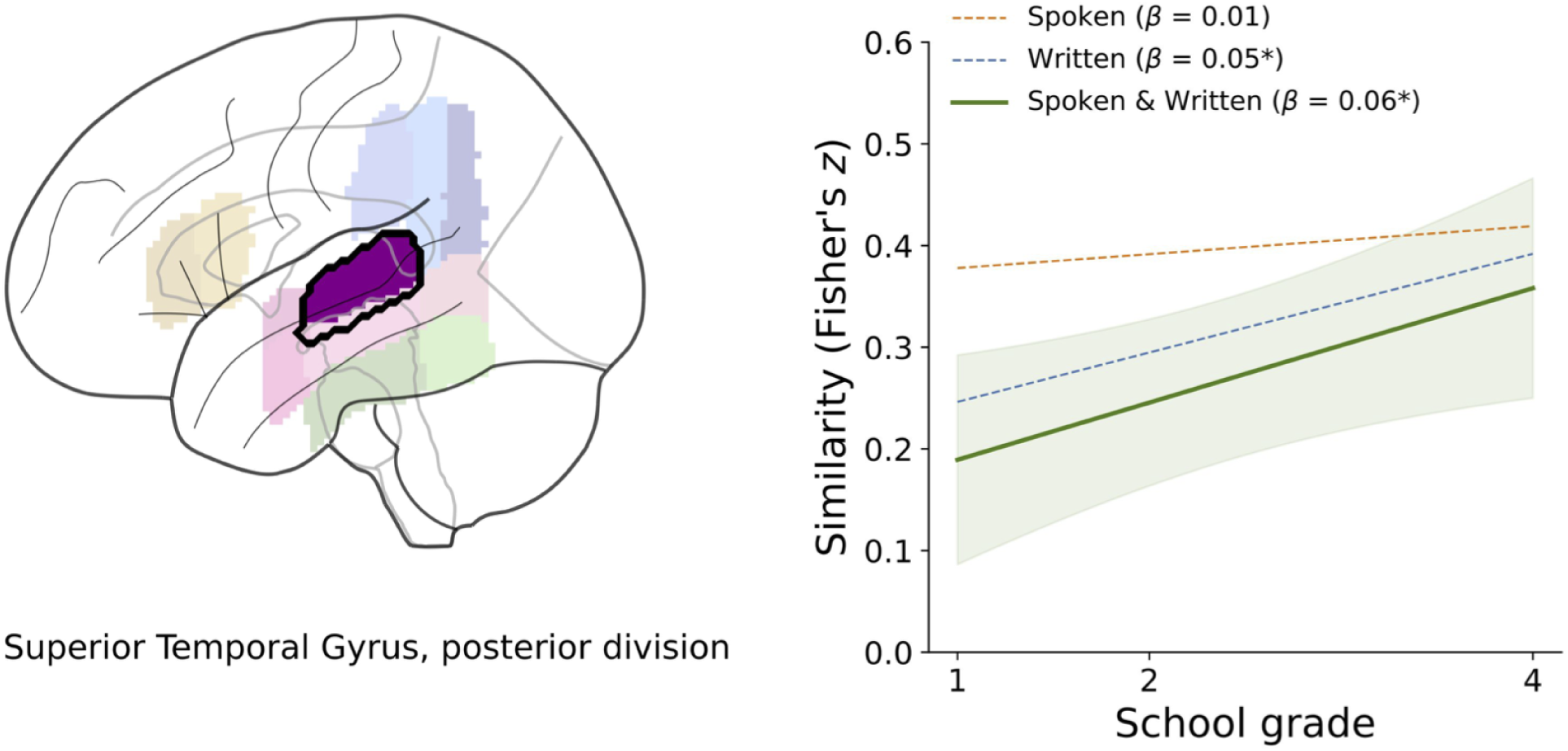
Cross-modal and unimodal similarity in the left posterior superior temporal gyrus. The solid line indicates cross-modal similarity and the dashed lines indicate unimodal similarity across school grades. Shaded areas represent 95% confidence intervals for the estimated cross-modal audiovisual similarity. Asterisks mark statistically significant effects of school grade at a significance level at *p_FDR_* < .05.

Spoken, spoken words only (unimodal auditory similarity). Written, written words only (unimodal visual similarity). Spoken & Written, both spoken and written words (cross-modal audiovisual similarity. ^a^ROI, region of interest; opIFG, inferior frontal gyrus, pars opercularis; aSMG, supramarginal gyrus, anterior division; pSMG, supramarginal gyrus, posterior division; pSTG, superior temporal gyrus, posterior division; toMTG, middle temporal gyrus, temporooccipital part; TOFC, temporal occipital fusiform cortex. ^b^*Mean similarity (Fisher’s z)* (G1 / G2 / G4), mean z-transformed Pearson correlation coefficients for first, second, and fourth grade, respectively. Significance level: *pFDR* < .05. ^c^*SD* (G1 / G2 / G4), standard deviation of z-transformed Pearson correlation coefficients for first, second, and fourth grade, respectively. ^d^*β*, regression coefficient for school grade. ^e^*R^2^*, proportion of variance explained by the corresponding model.

## Discussion

In this study, we investigated whether the multivariate similarity of hemodynamic responses to spoken and written words changes when learning to read in primary school. To this end, we acquired and analyzed fMRI data of children learning the alpha-syllabic Devanagari script in the first, second, or fourth grade. Cross-modal audiovisual similarity increased significantly across school grades in left inferior frontal, inferior parietal, superior and middle temporal, and temporo-occipital cortices. These results were largely comparable to the effects found for unimodal visual similarity while no significant effects were obtained for unimodal auditory similarity.

Significant differences in audiovisual similarity between lower and higher primary school grades were observed not only in the left pSTG, as hypothesized, but throughout the language system, in line with a recent meta-analysis of reading-related brain areas (Turker et al., 2025). Accordingly, we found evidence for a developmental trajectory, during which hemodynamic responses during visual word recognition become gradually aligned with those for auditory word comprehension. This finding provides further empirical support for the neuronal recycling hypothesis which posits that pre-existing neural resources evolved for spoken language processing are reused for written language processing by means of enculturation (Dehaene & Cohen, 2007; Dehaene et al., 2010; Rueckl et al., 2015). Moreover, our multivariate results complement prior univariate work showing overlapping activity during auditory and visual language processing in left-hemispheric perisylvian language areas by suggesting that this kind of processing leads to an alignment of activity patterns within these areas (Chyl et al., 2018; Dębska et al., 2021; Dębska et al., 2026; McNorgan et al., 2014; Preston et al., 2016).

The current study adds a developmental perspective to previous adult studies focusing on the abugida-based alphasyllabic Devanagari script, which is one of the most widely used writing systems in the world, and mastered by about half a billion readers (Hervais-Adelman et al., 2022; Skeide et al., 2017). Compared to alphabetic scripts, in which children learn to map graphemes on phonemes and vice versa, Devanagari requires mapping visually more complex characters (abugidas) on the corresponding consonant-vowel combinations that resemble syllable-like structures. Accordingly, given this visual and phonological complexity, efficient word recognition in Devanagari might require more intensive practice than in alphabetic scripts (Dehaene et al., 2015). This cultural difference might be especially relevant to alphabetic scripts such as Spanish or German in which grapheme-to-phoneme mapping is so transparent that word recognition typically emerges by the end of the first school year (Ziegler & Goswami, 2006). It can thus be expected that the similarity of spoken and written word representations will differ significantly between first and second graders learning a transparent alphabetic script. This effect was not found in the current Devanagari sample, in which particularly fourth graders revealed more similar representations. Comparative follow-up studies are required to test this prediction.

One limitation of the current study is that we cannot rule out that the observed increase in audiovisual similarity could be driven by cognitive differences extending beyond reading experience, particularly attention and working memory skills related to school grades. Moreover, the cross-sectional design of the present study precludes tracking developmental changes within individuals. A remaining challenge for future studies is therefore to track the same children longitudinally from first to fourth grade. An additional limitation is that, in contrast to previous studies (McNorgan et al., 2014; Preston et al., 2016; Dehaene-Lambertz et al., 2018), we did not collect standardized behavioral assessment of language skills related to reading performance outside the scanner. The decision not to collect behavioral data was made a priori, given that prior research has demonstrated that a sample size of several hundred or even thousand datasets is required to detect robust brain-behavior associations (Marek et al., 2022; Tervo-Clemmens et al., 2023).

## Conclusion

Leveraging a multivariate analysis of hemodynamic responses during word recognition in Devanagari, we demonstrated that brain responses to written words gradually align with brain responses to spoken words in perisylvian language areas when learning to read in primary school. These findings suggest that the language system becomes gradually more responsive to information from both sensory modalities in the course of formal instruction.

## Methods

### Participants

We recruited a total of 93 children from primary schools in New Delhi, India: first grade (*n* = 30), second grade (*n* = 27), and fourth grade (*n* = 35). All methods were carried out in accordance with the Declaration of Helsinki. All experimental protocols were approved by the Mahajan Imaging Ethics Committee in New Delhi, India (11/10/2018), the Ethics Committee of the Medical Faculty of the University of Leipzig, Germany (457/19-ek) and the Ethics Review Unit of the European Research Council Executive Agency (Ref. Ares(2020)307726 - 17/01/2020). Informed consent was obtained from a parent, and the children gave documented verbal assent to participate in the study. The rationale for investigating children learning the alpha-syllabic, abugida-based Devanagari script was that this writing system is understudied relative to alphabetic scripts like English (Kumar et al., 2010). All participants met the following inclusion criteria that were confirmed by parental reports: (1) no contraindications for MRI scanning (e.g., absence of metallic implants or relevant medication), (2) no hearing or vision impairments, (3) no diagnosis of neurological and neurodevelopmental disorders (including language disorders). Of the 93 participants initially recruited, 32 were excluded based on the following criteria: (1) excessive head motion, defined as framewise displacement exceeding 2.4 mm in more than 10% of fMRI volumes, (2) insufficient engagement in the semantic judgment task, defined as a response rate below 50% within a run, or (3) having fewer than three available runs, corresponding to less than 50% of all stimulus events. The final dataset therefore included 61 children: *n* = 18 first graders, *n* = 19 second graders, and *n* = 24 fourth graders. There were no significant associations between school grades and the number of available runs (*β* = 0.21, *t* = 1.53, *p* = 0.12), task response rate (*β* = 0.87, *t* = 1.60, *p* = 0.10) or task accuracy (*β* = 0.98, *t* = 1.31, *p* = 0.19). Reaction time and head motion (frame-wise displacement), however, decreased significantly across school grades (reaction time: *β* = -0.07, *t* = -3.56, *p* < 0.05; frame-wise displacement: *β* = -0.03, *t* = -3.37, *p* < 0.05) and were thus included as covariates of no interest in the statistical models.

### MRI data acquisition

MRI data were acquired on a GE SIGMA Architect 3T scanner with a 48 channel head coil. T1-weighted MPRAGE images were recorded with the following parameters: TE = 3.12 ms, TR = 2,100 ms, TI = 900 ms, flip angle = 8°, field of view = 256 × 256 mm, matrix size = 256 × 256 voxels, slice thickness = 1 mm, no gap between slices, voxel size = 1 × 1 × 1 mm, slice orientation = axial, and phase encoding direction = right-left. T2*-weighted EPI images (130 fMRI volumes per run in six total runs) were recorded with the following parameters: TE = 30 ms, TR = 2,000 ms, flip angle = 90°, number of volumes = 130, field of view = 192 × 192 mm, in plane matrix size = 64 × 64 voxels, slice thickness = 3 mm, inter-slice gap = 0.2 mm, voxel size = 3 × 3 × 3 mm, slice orientation = axial, phase encoding direction = anterior-posterior, slice order = interleaved ascending, and multiband acceleration factor = 2.

### MRI data preprocessing

Preprocessing was performed using *fMRIPrep* 25.1.4, which is based on *Nipype* 1.10.0.

### Anatomical data preprocessing

A single T1-weighted (T1w) image is found within the input BIDS dataset. The T1w image is corrected for intensity non-uniformity (INU) with `N4BiasFieldCorrection` (Tustison et al. 2010), distributed with ANTs 2.6.2 (Avants et al. 2008, RRID:SCR_004757), and use as T1w-reference throughout the workflow. The T1w-reference is then skull-stripped with a *Nipype* implementation of the `antsBrainExtraction.sh` workflow (from ANTs), using OASIS30ANTs as target template. Brain tissue segmentation of cerebrospinal fluid (CSF), white-matter (WM) and gray-matter (GM) is performed on the brain-extracted T1w using `fast` (FSL, RRID:SCR_002823, Zhang et al., 2001). Brain surfaces are reconstructed using `recon-all` (FreeSurfer 7.3.2, RRID:SCR_001847, Dale et al., 1999), and the brain mask estimated previously is refined with a custom variation of the method to reconcile ANTs-derived and FreeSurfer-derived segmentations of the cortical gray-matter of Mindboggle (RRID:SCR_002438, Klein et al., 2017). Volume-based spatial normalization to two standard spaces (MNI152NLin2009cAsym, MNIPediatricAsym:cohort-4) is performed through nonlinear registration with `antsRegistration` (ANTs 2.6.2), using brain-extracted versions of both T1w reference and the T1w template. The following templates are selected for spatial normalization and accessed with *TemplateFlow* (24.2.2, Ciric et al., 2022) and included:

- *ICBM 152 Nonlinear Asymmetrical template version 2009c* (Fonov et al., 2009; RRID:SCR_008796; TemplateFlow ID: MNI152NLin2009cAsym)
- *MNI Pediatric Asymmetrical template for ages 7.5-13.5 years* (RRID:SCR_008796; TemplateFlow ID: MNIPediatricAsym:cohort-4).

### Functional data preprocessing

For each of the 3-8 BOLD runs found per subject (across all sessions), the following preprocessing is performed. First, a reference volume was generated, using a custom methodology of *fMRIPrep*, for use in head motion correction. Head-motion parameters with respect to the BOLD reference (transformation matrices, and six corresponding rotation and translation parameters) are estimated before any spatiotemporal filtering using `mcflirt` (FSL, Jenkinson et al., 2002). The BOLD reference is then co-registered to the T1w reference using `bbregister` (FreeSurfer) which implements boundary-based registration (Greve & Fischl, 2009). Co-registration is configured with six degrees of freedom. Several confounding time-series are calculated based on the *preprocessed BOLD*: framewise displacement (FD), DVARS and three region-wise global signals. FD was computed using two formulations following Power (absolute sum of relative motions, Power et al., 2014) and Jenkinson (relative root mean square displacement between affines, Jenkinson et al., 2002). FD and DVARS are calculated for each functional run, both using their implementations in *Nipype* (following the definitions by Power et al., 2014). The three global signals are extracted within the CSF, the WM, and the whole-brain masks. Additionally, a set of physiological regressors are extracted to allow for component-based noise correction (*CompCor*, Behzadi et al., 2007). Principal components are estimated after high-pass filtering the *preprocessed BOLD* time-series (using a discrete cosine filter with 128s cut-off) for the two *CompCor* variants: temporal (tCompCor) and anatomical (aCompCor). tCompCor components are then calculated from the top 2% variable voxels within the brain mask. For aCompCor, three probabilistic masks (CSF, WM and combined CSF+WM) are generated in anatomical space. The implementation differs from that of Behzadi et al. in that instead of eroding the masks by 2 pixels on BOLD space, a mask of pixels that likely contain a volume fraction of GM is subtracted from the aCompCor masks. This mask is obtained by dilating a GM mask extracted from the FreeSurfer’s *aseg* segmentation, and it ensures components are not extracted from voxels containing a minimal fraction of GM. Finally, these masks are resampled into BOLD space and binarized by thresholding at 0.99 (as in the original implementation). Components are also calculated separately within the WM and CSF masks. For each CompCor decomposition, the k components with the largest singular values are retained, such that the retained components’ time series are sufficient to explain 50 percent of variance across the nuisance mask (CSF, WM, combined, or temporal). The remaining components are dropped from consideration. The head-motion estimates calculated in the correction step are also placed within the corresponding confounds file. The confound time series derived from head motion estimates and global signals are expanded with the inclusion of temporal derivatives and quadratic terms for each (Satterthwaite et al., 2013). Frames that exceeded a threshold of 0.5 mm FD or 1.5 standardized DVARS are annotated as motion outliers. Additional nuisance time series are calculated by means of principal components analysis of the signal found within a thin band (*crown*) of voxels around the edge of the brain, as proposed by (Patriat et al., 2017). The BOLD time-series are resampled onto the following surfaces (FreeSurfer reconstruction nomenclature): *fsnative*, *fsaverage5*. All resamplings can be performed with a single interpolation step by composing all the pertinent transformations (i.e. head-motion transform matrices, susceptibility distortion correction when available, and co-registrations to anatomical and output spaces). Gridded (volumetric) resamplings are performed using nitransforms, configured with cubic B-spline interpolation. Non-gridded (surface) resamplings are performed using mri_vol2surf (FreeSurfer).

Many internal operations of *fMRIPrep* use *Nilearn* 0.11.1 (Abraham et al., 2014, RRID:SCR_001362), mostly within the functional processing workflow. For more details of the pipeline, see the section corresponding to workflows in *fMRIPrep*’s documentation (https://fmriprep.org/en/latest/workflows.html).

Copyright Waiver: The above boilerplate text is automatically generated by *fMRIPrep* with the express intention that users should copy and paste this text into their manuscripts *unchanged*. It is released under the CC0 license.

### Experimental paradigm

During fMRI scanning, stimuli were presented in four conditions: written words, written pseudowords, spoken words, and spoken pseudowords. All Hindi words were animal names that were all masculine nouns containing one or two syllables and three to six phonemes. Pseudowords were generated by substituting the initial consonant and vowel of each word with phonemes sharing the same articulatory place. Visual stimuli were presented in the middle of the screen in black font on a white background. The screen was positioned behind the scanner bore and viewed through a mirror mounted on the head coil. Auditory stimuli were recorded by a male native Hindi speaker. The experiment comprised up to eight runs and employed an event-related design. Each run consisted of 36 pseudo-randomized trials balanced across four conditions. In each trial, a stimulus was presented for 3 seconds, followed by a baseline period during which a black fixation cross was displayed for a pseudo-random duration ranging from 3 to 5 seconds. During spoken conditions, a black fixation cross remained visible at the center of the screen. Each run lasted approximately 4 minutes and 30 seconds, and the total experimental duration was approximately 42 minutes. To keep participants engaged, we employed a semantic judgement task, and instructed them to indicate whether a stimulus represented an animal or not, using an MR-compatible button box. The experiment was programmed and presented using PsychoPy (v2021; Peirce et al., 2019).

### Subject-level fMRI analysis

We estimated a general linear model (GLM) implemented in Nilearn (v0.12.1) for Python (v3.12.12). Separately for each participant and each run, fMRI time series were standardized to percent signal change and spatially smoothed using a 5 mm full width at half maximum (FWHM) Gaussian kernel. This smoothing size aligns with prior developmental fMRI literature using kernels ranging from 5 to 10 mm to examine the similarity of hemodynamic activation (Chyl et al., 2021; Cohen et al., 2019; Mathur et al., 2020). Stimulus events were convolved with the canonical hemodynamic response function implemented in SPM to obtain predicted blood-oxygen-level dependent (BOLD) signals. We estimated four contrasts, including written words (written words > baseline), written semantic information (written words > pseudowords), spoken words (spoken words > baseline), and spoken semantic information (spoken words > pseudowords). To account for motion- and physiology-related noise, several nuisance regressors were included in the design matrix: (1) six head motion parameters (translations and rotations in three directions) estimated by fMRIPrep, (2) the top six anatomical CompCor components estimated by fMRIPrep, (3) cosine regressors to high-pass filter the data with a cutoff frequency of 0.01 Hz (approximately 128 seconds), (4) spike regressors modeling non-steady-state volumes identified at the beginning of each run estimated by fMRIPrep, (5) spike regressors modeling volumes with excessive motion, defined as framewise displacement greater than 0.5 mm estimated by fMRIPrep. The run-level contrasts yielded maps of effect size (*β)*, variance, *t*-statistics, and *z*-scores. We used the resulting *β*-maps for subsequent cross-modal and unimodal similarity estimation.

### Regions of Interest

We selected twelve cortical masks in the left hemisphere as regions of interest (ROIs) using the Harvard–Oxford cortical atlas (cort-maxprob-thr25-2mm) provided in Nilearn. ROI masks were transformed into the pediatric template space using antsRegistration (v0.6.1 ANTsPy). Following Karipidis et al. (2021), the transformed ROI masks were restricted to voxels with a gray matter probability > 0.5 according to the tissue probability map of the pediatric MNI template. Following a number of previous developmental language processing studies, we selected this atlas as it delineates subregions within canonical language areas, particularly in Broca’s area, including the pars opercularis and pars triangularis of the inferior frontal gyrus (Youssofzadeh et al., 2018; Krishnan et al., 2021; Li et al., 2017; Nuñez et al., 2011). We adopted a ROI-based approach rather than an exploratory whole-brain approach because canonical written and spoken language areas have already been identified in large-scale meta-analyses comprising over 400 studies (Malik-Moraleda et al., 2022; Rueckl et al., 2015; Turker et al., 2023; Turker et al., 2025).

### Cross-modal and unimodal similarity estimation

To estimate cross-modal and unimodal similarity, we performed a correlation analysis within the ROIs, adopting the approach used in prior cross-modal language studies (Chyl et al., 2021; Marks et al., 2019; Rueckl et al., 2015). Specifically, similarity was calculated by averaging Fisher *z*-transformed Pearson’s correlation coefficients across all pairwise comparisons of effect maps for spoken and written words across different runs. Comparisons within the same run were excluded to minimize confounding by temporally autocorrelated noise sources, including physiological fluctuations (e.g., respiratory and cardiac cycles) and scanner drift (e.g., slow fluctuations of the static magnetic field). In addition to cross-modal similarity, unimodal similarity was also computed following the same procedure but with pairwise comparisons across different runs corresponding to the same (auditory or visual) modality. Additionally, we also calculated similarity based on effect maps contrasting words against pseudowords to explore cross-modal semantic similarity.

### Statistical analyses

To compute main effects of school grade and ROI as well as the grade x ROI interaction effect, we conducted two-way mixed ANOVAs using the Pingouing python package (Vallat, 2018). Whenever the assumption of sphericity was violated, degrees of freedom were corrected using the Greenhouse–Geisser approach. Following the observation of a significant interaction effect, we tested for significant differences across school grades in each ROI by implementing post-hoc pairwise comparisons using Welch’s *t*-test to account for the unequal sample sizes across school grades (Zimmerman, 2004). The resulting 36 uncorrected *p*-values, corresponding to 3 comparisons × 12 ROIs, were adjusted using the false discovery rate (FDR) procedure (Benjamini–Hochberg method), based on a significance threshold of *q* < .05. To evaluate whether similarity increased or decreased with school grade, we fitted multiple linear regression models for each ROI, with school grade as the independent variable, similarity as the dependent variable, and sex, mean framewise displacement (head motion), and mean reaction times in the semantic judgement task as covariates of no interest. Similarity values with a Cook’s distance greater than 4/*n* = 0.066 (*n* = 61) were considered influential outliers and excluded from the regression models (Altman & Krzywinski, 2016).

## Acknowledgements

The authors would like to thank Ira Sarma and Siddharta Kotwal for their help with creating the stimulus material, Alexander Enge for his methodological advice on the experimental design, and Kabir Mahajan, Vidur Mahajan, Nisha Nasser, Madhuri Barnwal, Manmeet Singh, Nitesh Shisodia, Shivani Rastogi, Devanshi Khanna, Priyanka Singh, Kriti Tyagi, Annika Dold, Martin Nielebock, Lena Traser, Jasmin Wend, and Michael Vollmann for supporting data collection.

## Author contributions

K.M.: methodology, software, formal analysis, visualization, writing—original draft, writing—review and editing.

R.K.: methodology, writing—review and editing.

F.H.: writing—review and editing.

M.A.S.: conceptualization, data curation, methodology, resources, project administration, supervision, writing—review and editing.

## Data availability

Preprocessed data supporting the findings of this study are available from the corresponding author upon reasonable request and will be uploaded to an open repository after publication.

## Code availability

The code for all analyses is available at https://github.com/SkeideLab/SLANG-CROSS-analysis.

