## Supplementary Material for "Multivariate pattern similarity of hemodynamic brain responses to spoken and written words increases during primary school"

### Supplementary Methods

#### Audiovisual coactivation analysis

To investigate the relationship between similarity and overlap of hemodynamic responses elicited by spoken and written words, we additionally performed a conjunction analysis serving as an index of speech–print coactivation. Following previous studies, subject-level effect maps were smoothed using an 8 mm full-width at half-maximum isotropic Gaussian kernel. Audiovisual coactivation was defined at the single-subject level by identifying voxels displaying significant activation ($p < .05$) in the spoken and written word conditions, corresponding to a conjoint probability of $p < .0025$ (Dębska et al., 2021; Preston et al., 2016). In line with our cross-modal similarity analyses, the extent of coactivation within each region of interest (ROI) was quantified by calculating the mean number of coactivated voxels across all pairwise cross-modal comparisons between spoken and written words, excluding within-run comparisons. Subsequently, Spearman's rank correlation analyses were conducted to examine the relationship between audiovisual similarity and audiovisual coactivation for each ROI. The resulting *p*-values across the 12 ROIs were adjusted using the false discovery rate (FDR) procedure, with a significance threshold of $q < .05$.

### Supplementary Results

#### Relationship between audiovisual coactivation and multivariate similarity

We examined whether the extent of audiovisual coactivation was associated with cross-modal audiovisual similarity. Spearman's rank correlation analyses revealed a significant positive relationship between the number of coactivated voxels and audiovisual similarity across 9 out of 12 ROIs after FDR correction ($q$ < .05), including the left posterior superior temporal gyrus (pSTG) (Spearman $r(59)$ = 0.62, $p_{FDR}$ < .05). In contrast, no significant correlations were found in the anterior dvision of the left middle temporal gyrus (aMTG), the posterior division of the middle temporal gyrus (pMTG), and the posterior division of the temporal fusiform cortex (aTFC) (Fig. S1).

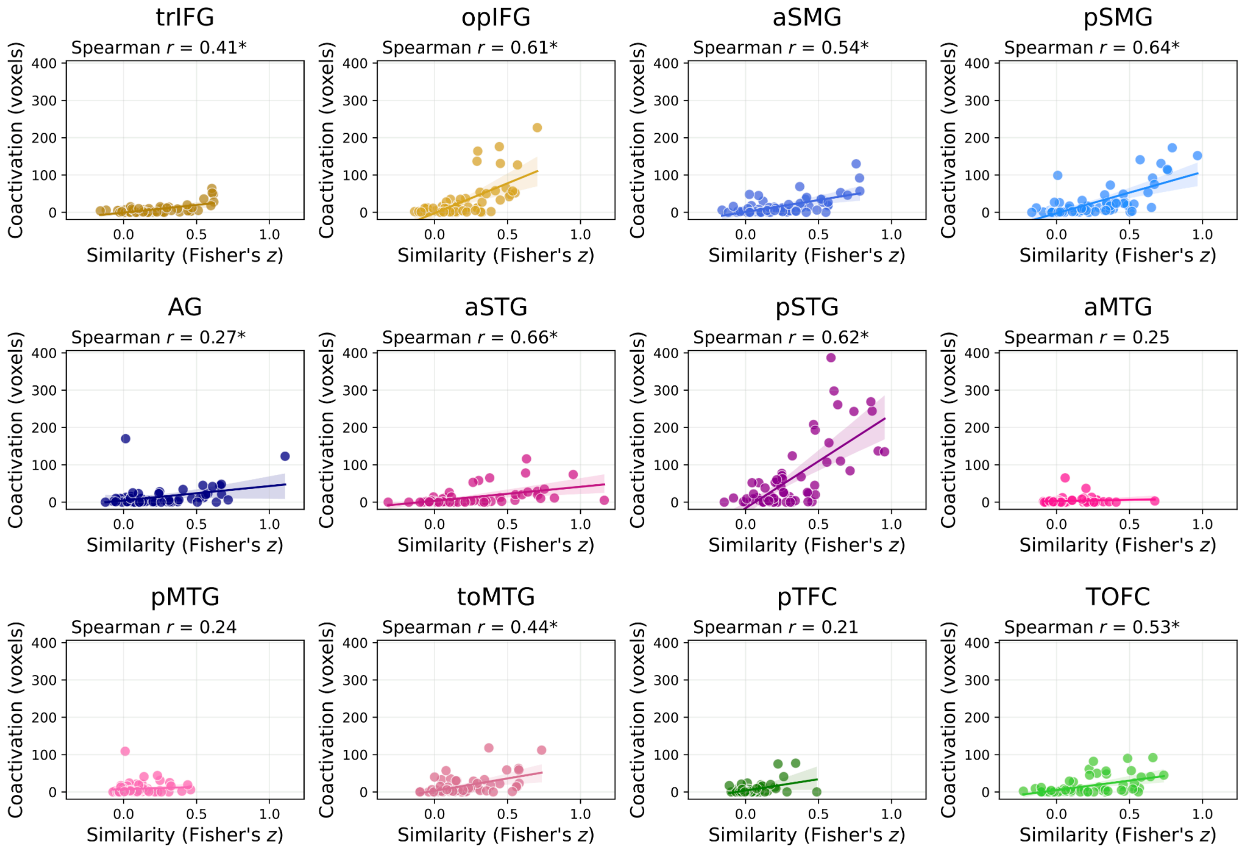

**Figure S1. Associations between audiovisual coactivation and multivariate similarity.** Scatter plots show the relationship between audiovisual similarity (Fisher's *z*, x-axis) and audiovisual coactivation (number of coactivated voxels, y-axis) across 12 ROIs. Each dot represents the value of an individual participant. Solid lines represent linear regression fits with 95% confidence intervals. Spearman's rank correlation coefficients are reported (Spearman's *r*). Asterisks (*) indicate significant correlations after FDR correction ($q < .05$). trIFG, inferior frontal gyrus, pars triangularis; opIFG, inferior frontal gyrus, pars opercularis; aSMG, supramarginal gyrus, anterior division; pSMG, supramarginal gyrus, posterior division; AG, angular gyrus; aSTG, superior temporal gyrus, anterior division; pSTG, superior temporal gyrus, posterior division; aMTG, middle temporal gyrus, anterior division; pMTG, middle temporal gyrus, posterior division; toMTG, middle temporal gyrus, temporooccipital part; pTFC, temporal fusiform cortex, posterior division; TOFC, temporal occipital fusiform cortex.
